# Short-term CaMKII inhibition with tatCN19o does not erase pre-formed memory and is neuroprotective in non-rodents

**DOI:** 10.1101/2023.01.23.523316

**Authors:** Nicole L. Rumian, Carolyn Nicole Brown, Tara B. Hendry-Hofer, Thomas Rossetti, James E. Orfila, Jonathan E. Tullis, Linda P. Dwoskin, Olivia R. Buonarati, John E. Lisman, Nidia Quillinan, Paco S. Herson, Vikhyat S. Bebarta, K. Ulrich Bayer

## Abstract

The Ca^2+^/calmodulin-dependent protein kinase II (CaMKII) is a central regulator of learning and memory, which poses a problem for targeting it therapeutically. Indeed, our study supports prior conclusions that long-term interference with CaMKII signaling can erase pre-formed memories. By contrast, short-term pharmacological CaMKII inhibition with tatCN19o interfered with learning in mice only mildly and transiently (for less than 1 h) and did not at all reverse pre-formed memories. This was at ≥500fold of the dose that protected hippocampal neurons from cell death after a highly clinically relevant pig model of transient global cerebral ischemia: ventricular fibrillation followed by advanced life support and electrical defibrillation to induce return of spontaneous circulation. Of additional importance for therapeutic development, cardiovascular safety studies in mice and pig did not indicate any concerns with acute tatCN19o injection. Taken together, even though prolonged interference with CaMKII signaling can erase memory, acute short-term CaMKII inhibition with tatCN19o did not cause such retrograde amnesia that would pose a contraindication for therapy.

## INTRODUCTION

Learning and memory are thought to require forms of synaptic plasticity, specifically including hippocampal long-term potentiation and depression (LTP and LTD) (1–3). LTP, LTD, and learning all require the Ca^2+^/calmodulin-dependent protein kinase II (CaMKII) and its autophosphorylation at T286 that generates Ca^2+^-independent “autonomous” kinase activity (4–6). Normal LTD additionally requires the inhibitory CaMKII autophosphorylation at T305/306 (7), whereas normal LTP instead requires the regulated CaMKII binding to the NMDA-type glutamate receptor (NMDAR) subunit GluN2B (8–10). In addition to LTP induction, CaMKII is thought to mediate LTP maintenance, although the mechanisms are less clear. Initially, LTP maintenance had been proposed to involve T286 autophosphorylation, but several lines of evidence argue for GluN2B binding as the more likely mechanism (6,11–13). The maintenance of LTP is also proposed to mediate maintenance of memory. However, to date, there is only one convincing study that directly indicated CaMKII involvement in memory maintenance in a hippocampus-dependent task (14). Notably, this study used transient overexpression of the CaMKII K42M mutant, which prevents nucleotide binding to the kinase domain and thereby interferes with both T286 autophosphorylation and GluN2B binding (15,16), i.e. both of the two potential CaMKII mechanisms proposed to function in LTP maintenance. Overexpression of the K42M mutant can affect endogenous wild type CaMKII by integrating together into holoenzymes and thereby acting as dominant negative. Using a viral vector that was based on the herpes simplex virus (HSV), the CaMKII K42M mutant was transiently expressed in hippocampus after training rats in a conditioned place avoidance task. This transient expression persistently erased the memory of the avoidance task (14). A similar HSV-based approach was used to show persistent reversal of maladaptive memory related to amphetamine addiction, in this case by transient viral expression of the K42M mutant in the nucleus accumbens shell (17). Both studies are strong, but both come with caveats (6): To claim memory erasure, it has to be demonstrated that the K42M mutant is indeed no longer expressed at the time of the memory test; otherwise, failure in the memory test could be attributed to acute disturbance of memory retrieval by the still expressed mutant, rather than to persistent erasure of the memory. In the addiction study, CaMKII expression level in the nucleus accumbens shell was back to baseline at the time of testing (17), however, as non-tagged CaMKII was used, it is formally possible that this was due to adaptive downregulation of endogenous wild type kinase rather than elimination of the K42M mutant. In the hippocampal memory study, the transiently expressed CaMKII was either directly GFP-tagged (for wildtype and T286D/T305/306AA) or was non-tagged but included a cassette for separate GFP expression on the same vector (for K42M). However, elimination of expression was shown only for the less informative CaMKII tool mutant, the constitutively active T286D/T305/306AA mutant (14). Thus, in both cases, K42M expression may have persisted on the test day, if the K42M mutant is more stable or more highly expressed than the CaMKII T286D/T305/306AA mutant.

Here we found that both CaMKII wildtype and the K42M mutant are indeed significantly higher expressed in HEK cells than T286D/T305/306AA. However, after HSV-mediated expression in rat hippocampus, GFP expression was no longer detected on the test day for either mutant, thus supporting the original conclusion that long-term interference with CaMKII signaling can erase memories. By contrast, we found that short-term CaMKII inhibition with the neuroprotective peptide tatCN19o (18,19) transiently interfered with learning, but did not reverse pre-formed memories, even at ≥500fold of a neuroprotective dose determined previously in mice and here in pig. Thus, short-term inhibition of CaMKII with tatCN19o does not cause retrograde amnesia. Together with the demonstrated efficacy also in a non-rodent species, these findings indicate feasibility of short-term CaMKII inhibition as neuroprotective therapy for cerebral ischemia.

## RESULTS

### The CaMKII tool mutants show different expression levels in HEK293 cells

To compare the expression levels of CaMKII wildtype and mutants, we used a bi-cistronic expression vector in which an internal ribosomal entry site (IRES) allows expression of two different proteins from the same mRNA transcript: YFP-CaMKII and mTurquoise (Figure 1A). Twenty-four hours after transfection, HEK293 cells were imaged for expression analysis by ratiometric quantification of the YFP-CaMKII fluorescence relative to the mTurquoise fluorescence (Figure 1B). Significantly higher levels of expression were observed for CaMKII wildtype and the K42M mutant compared to the T286D/T305/306AA mutant, although the size of the effect was relatively small (Figure 1C).

**Figure 1:**
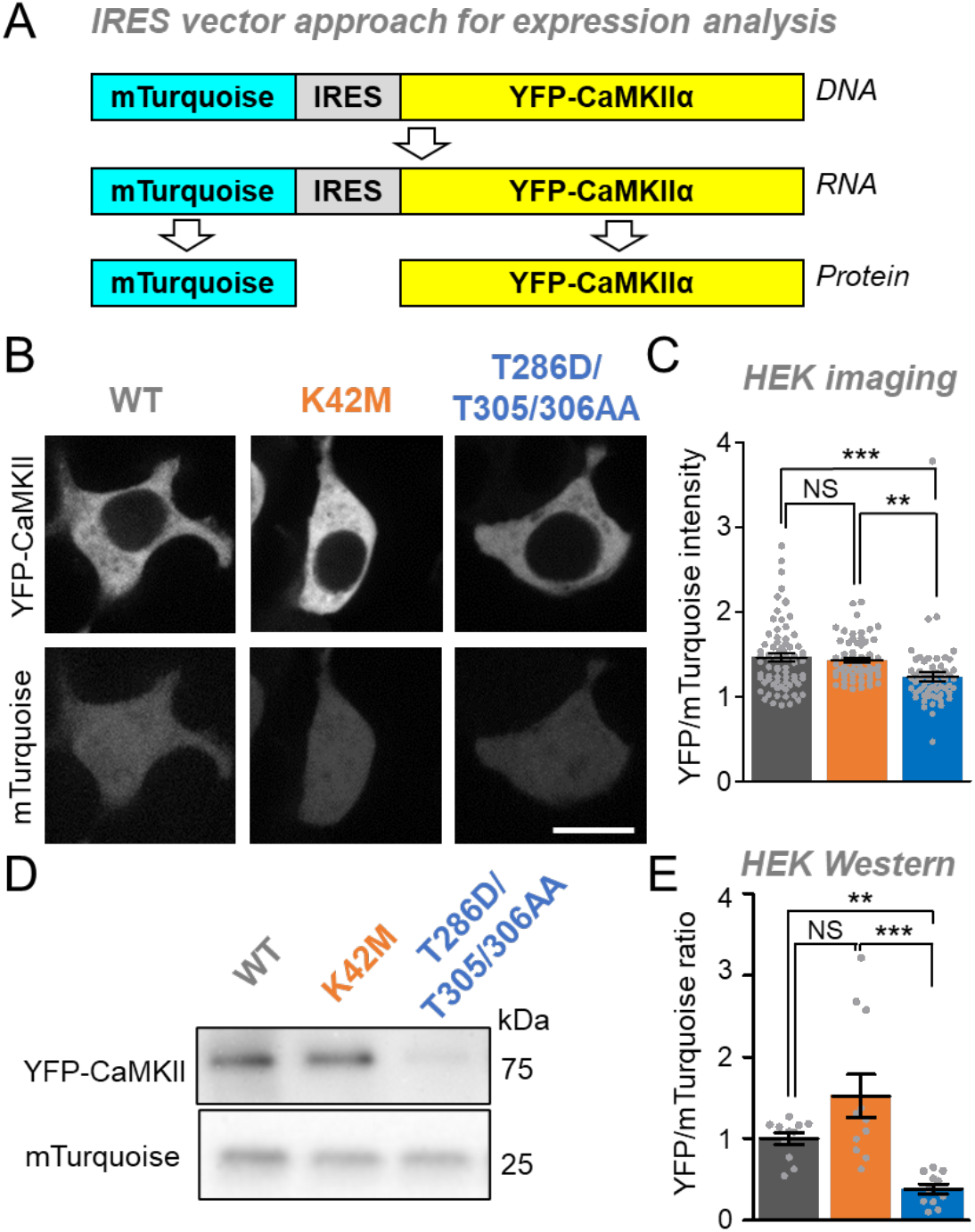
Differential expression levels of CaMKII tool mutants in HEK293 cells.

Error bars indicate SEM in all panels. **P<0.01; ***P<0.001; NS, not significant.

(A) An mTurquoise-IRES-YFP-CaMKII vector is expressed as a single mRNA transcript that undergoes translation separately for mTurquoise and YFP-CaMKII.

(B) Representative images of mTurquoise and YFP-CaMKII in HEK cells.

(C) Quantification of YFP-CaMKII wildtype (n=59), K42M (n=62), and T286D/T305/306AA (N=70) images corrected for mTurquoise in HEK cells.

(D) Representative images of blots for mTurquoise and YFP-CaMKII from HEK cell lysates (Scale bar = 10 μm).

(E) Quantification of mTurquoise and YFP-CaMKII wildtype (n=11), K42M (n=11), and T286D/T305/306AA (n=11) blots from HEK cell lysates.

Next, we used Western analysis for additional independent assessment of the expression levels of the CaMKII wildtype and the two mutants. YFP-CaMKII and mTurquoise were detected with the same anti-GFP antibody on the same blot, with distinction of the two proteins based on their different sizes (Figure 1D). Again, this allowed ratiometric quantification of the YFP-CaMKII expression, and again CaMKII wildtype and the K42M mutant were significantly more highly expressed compared to the T286D/T305/306AA mutant (Figure 1E). These results are consistent with the findings from the imaging analysis, however, the effect size that was detected by Western blot was much larger (compared Figure 1E and 1C). One possibility for the smaller apparent effect size detected by imaging was that YFP-CaMKII may be degraded in a way that still leaves the fluorophore intact, thereby leading to an underestimation of the degree of degradation; however, Western analysis did not find any indication for such a selective degradation. (Figure S1)

Taken together, in HEK293 cells, CaMKII wildtype and the K42M mutant are significantly more highly expressed compared to the T286D/T305/306AA mutant. This conclusion is supported by both of our approaches, even though the single cell analysis by imaging appeared to be less sensitive.

### The mechanism for lower expression of the T286D/T305/306AA mutant is unclear

The decreased expression level caused by the T286/T305/306AA mutation could be due to either decreased production or increased degradation. An especially attractive hypothesis was an increase in proteasome-mediated degradation, as the proteasome can be stimulated by CaMKII activity (20) and the T286D/T305/306AA mutant is constitutively active. In order to test this hypothesis, we set out to compare the effect of the proteasome inhibitor MG-132 (at 50 μM) on the relative expression levels of CaMKII wildtype and the two tool mutants. If the differences in expression levels are entirely due to difference in degradation by the proteasome, then inhibition of the proteasome should completely equalize these expression levels. However, our results remain inconclusive: Prolonged treatment with MG-132 was toxic to the cells in our hands. Treatment for 4 h did not normalize the expression level of the T286/T305/306AA mutant (Figure S2), but based on CaMKII half-life estimates of 1-3 days (21,22), normalization of expression would not be expected after such short time of proteasome inhibition.

### No evidence for differential expression levels in hippocampal neurons

Next, we decided to compare the expression of CaMKII wildtype and the two tool mutants in cultured hippocampal neurons (Figure 2A), as these neurons provide a more physiologically relevant environment. In the hippocampal neurons, any potential differences in the detected expression levels between CaMKII wildtype and the mutants were not statistically significant (Figure 2A,B). However, due to the low transfection efficiency in primary hippocampal cultures, the assessment of expression was restricted to single cell analysis by imaging, which was not as sensitive at detecting the different levels of expression in HEK cells as the Western analysis (see Figure 1).

**Figure 2:**
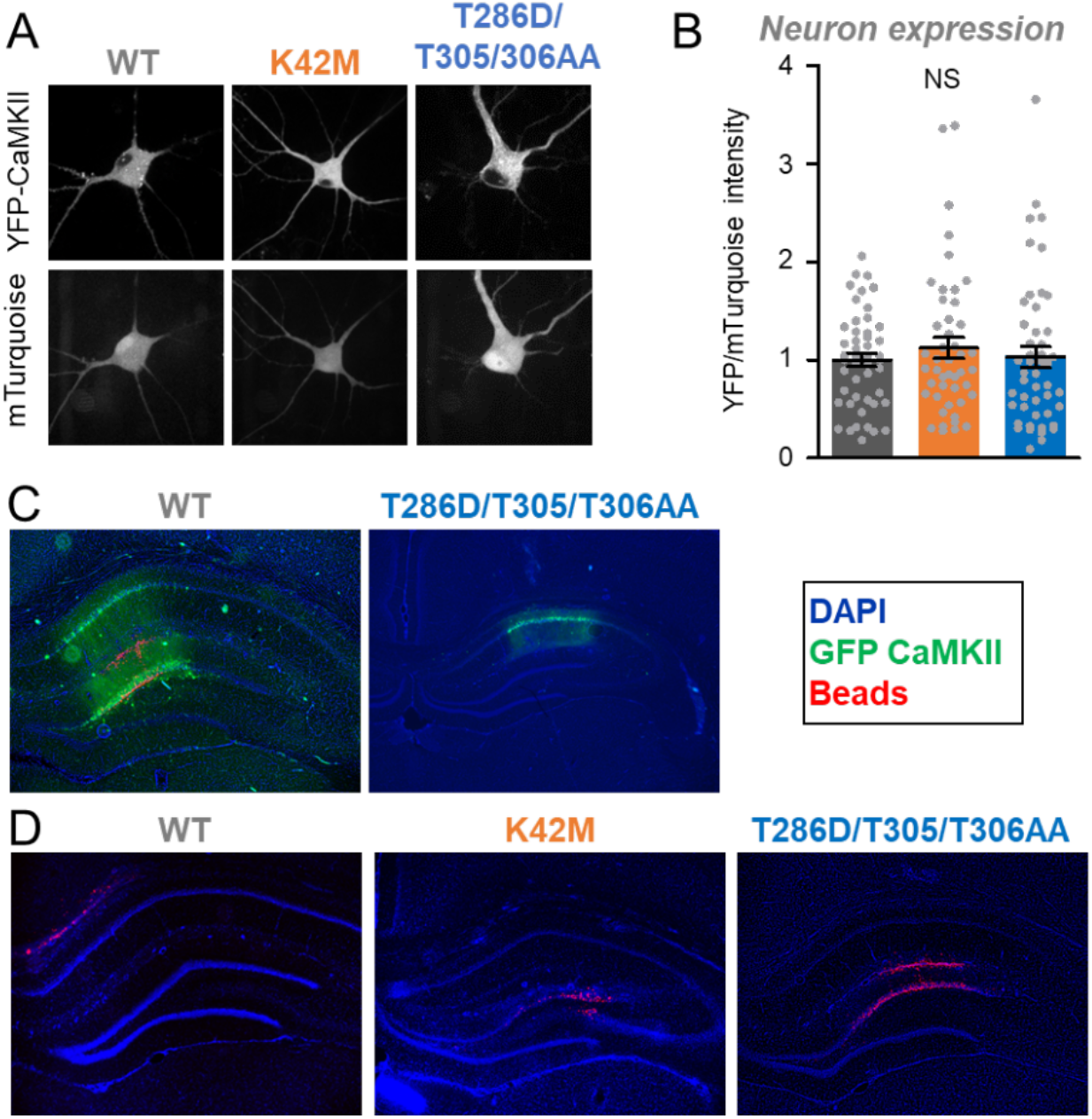
Expression of CaMKII tool mutants in hippocampal neurons in cultures and in vivo.

Error bars indicate SEM in all panels. NS, not significant.

(A) Representative images of mTurquoise and YFP-CaMKII in dissociated hippocampal neurons.

(B) Quantification of YFP-CaMKII wildtype (n=48), K42M (n=46), and T286D/T305/306AA (n=48) images corrected for mTurquoise in dissociated hippocampal neurons.

(C) wildtype and T286D/T305/T306AA expression on day 3 after HSV.

(D) No GFP expression on day 9 after HSV in example images for T286D/T305/306AA (n=6), wildtype (n=5), and K42M (n=6).

### The HSV-mediated transient expression of GFP-CaMKII in hippocampus *in vivo* is extinguished on day 9 after infection

For the GFP-CaMKII T286D/T305/306A mutant, a previous study showed that HSV-mediated expression of GFP in the hippocampus is strong on day 3 after infection but completely extinguished by day 9 after infection (14), and another example for this effect is also shown here (Figure 2C,D). Even though the original study did not report the expression levels for GFP-CaMKII wildtype or for the polycistronic construct for separate expression of GFP and the CaMKII K42M mutant, expression was analyzed for all animals after their final behavioral testing on day 9 after infection. On day 9, no GFP expression was detectable for any of the three constructs, with example images shown here (Figure 2D), confirming that all constructs expressed transiently and that GFP expression was extinguished on the day of behavioral testing for memory extinction in the original study. This indicates that the expression of the CaMKII K42M mutant was indeed transient and caused persistent interference with memory maintenance rather than acute interference with memory retrieval.

### CaMKII inhibition with tatNC19o interferes with learning only mildly and very transiently

The experiments that suggested a role of CaMKII in memory maintenance used a prolonged transient interference with CaMKII signaling over a period of multiple days, achieved by viral expression of the CaMKII K42M mutant (14,17). This raised the possibility that short-term pharmacological CaMKII inhibition could also affect memory, which would be a counter-indication for a promising neuroprotective treatment of cerebral ischemia with the CaMKII inhibitor tatCN19o (18,19). Thus, we decided to directly test the effect of tatCN19o on learning and memory in a hippocampus-dependent contextual fear conditioning paradigm (Figure 3A): During the learning session, mice are placed in a box where they receive a foot-shock. During the test session 24 h later, their memory is assessed by placing the mice back into the same box and measuring their freezing behavior without any shock. First, we tested the effect of tatCN19o on learning, by injecting it i.v. at various time points at a dose of 1 mg/kg. Notably, this is 100x of the dose that showed maximal neuroprotection after global cerebral ischemia (18). Only the injection at the shortest time point before the learning session (15 min) showed any apparent impairment in learning (Figure 3B,C). Any other timepoints of pre-learning injections (1 to 16 h) did not show any impairment at all. During the first 2 min of the test session, the cohort with 15 min pre-training injection was the only one with significantly less freezing behavior than the other groups (Figure 3B). During the last 4 min of the test session, the same cohort was also the only one with any statistically significant differences, but in this case only in comparison to the cohort with 4 h pre-training injection (Figure 3C). Thus, any effects of acute tatCN19o injection on learning appear to be relatively mild and are certainly very short-lived, and therefore do not pose a counter-indication for therapeutic applications.

**Figure 3:**
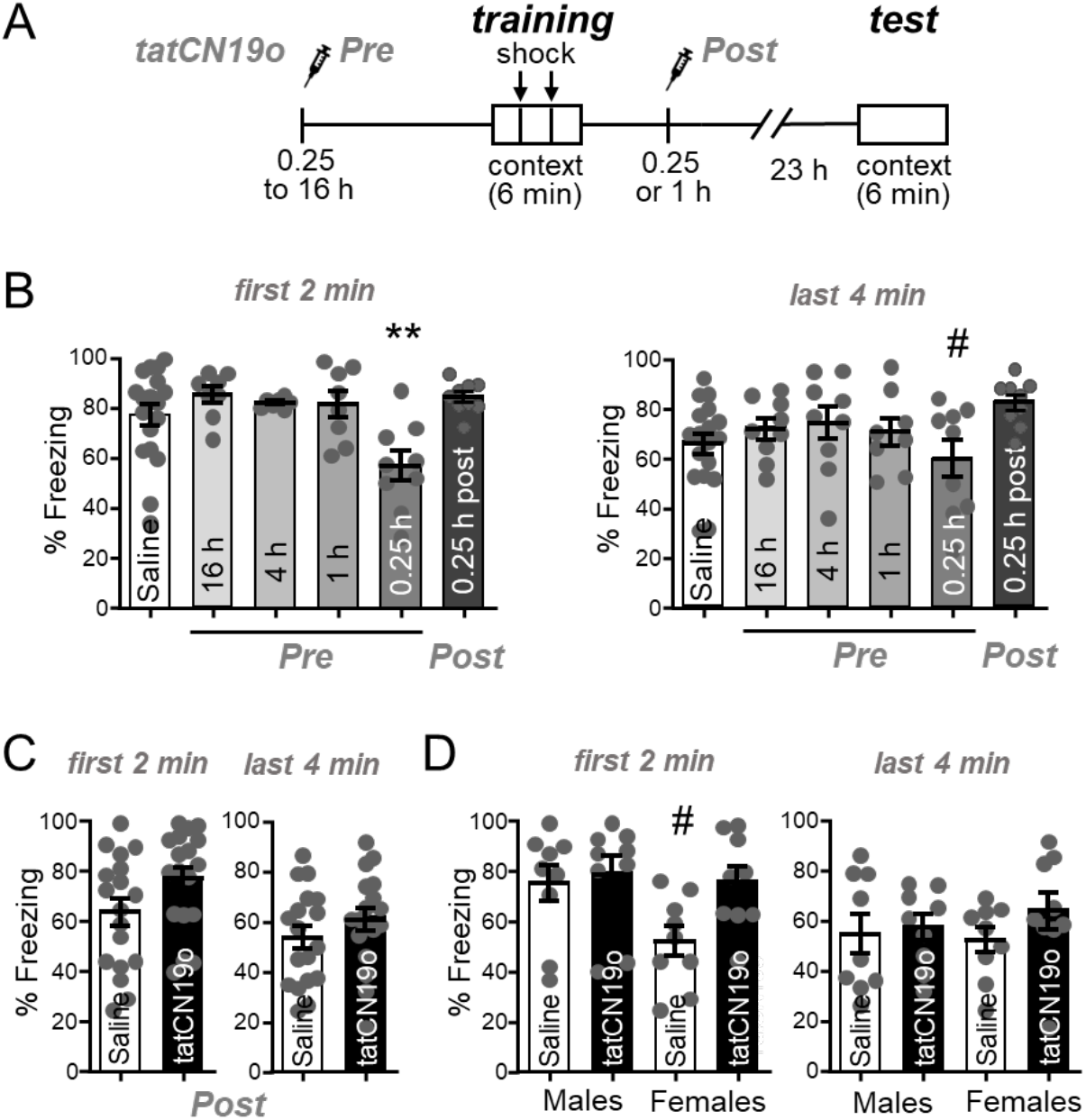
tatCN19o transiently affects learning but not memory.

Error bars indicate SEM in all panels.

(A) Schematic of the Contextual Fear Conditioning behavioral task (CFC) and timeline of saline or tatCN19o injections. Injections (1 mg/kg or 10 mg/kg) occurred either before (16, 4, 1, or 0.25 h) or after (0.25 h) the training session (2 foot shocks over the 6 min trial). For the test session 24 h later, the mice were returned to the chamber but received no foot shocks. Freezing response was measured across both days.

(B) Quantification of freezing behavior during the first 2 min or last 4 min of the test session of the first set of experiments. Only the 15 min tatCN19o injection pre-learning cohort demonstrated a significant reduction in freezing behavior during the first 2 min (*p<0.05 in one-way ANOVA with Bonferroni post-hoc test compared to the saline control condition, n=18, 8, 7, 8, 9, 9 animals) or the last 4 min (one-way ANOVA, n=18, 9, 9, 8, 9, 9 animals; # p<0.05 compared to the condition of 4 h pre-injection).

(C) Quantification of freezing behavior during the first 2 min and last 4 min of the second set of experiments, with all injections 1 h after learning (N.S. in unpaired t-test, n=18, 18 animals).

(D) Quantification of freezing behavior during the first 2 min and last 4 min of the second set of experiments, separated by sex. There was now statistical difference detected by two-way ANOVA (n=9, 9, 9, 9 animals). The apparent reduced freezing in the first 2 min period in females without compared to with tatCN19o treatment was significant only by direct comparison in an unpaired t-test (#: p<0.05) and would indicate enhanced, not decreased, memory after tatCN19o injection.

### CaMKII inhibition with tatCN19o does not interfere with preformed memory

In order to test possible effects of tatCN19o on pre-formed memories rather than on learning, we additionally injected tatCN19o after the learning session. In the first experiment, even injection of 10 mg/kg tatCN19o (i.e. 1,000x the therapeutic dose in cerebral ischemia) at 15 min after the learning session showed no apparent reduction in memory (Figure 3B,C). However, this first experiment was done in parallel with a learning experiment, and the control condition was a pre-learning saline injection, in contrast to the post-learning injection of tatCN19o that tests effects on memory. Thus, we performed an additional independent experiment that directly compared post-learning injection of 10 mg/kg tatCN19o versus saline at the same 1 h time point after the learning session (Figure 3D,E). In this experiment, both male and female mice were analyzed. Again, no effect of tatCN19o on memory was detected, neither when the sexes were analyzed together and compared by t-test (Figure 3D) nor when the sexes were separated and analyzed by two-way ANOVA (Figure 3E). The only apparent difference in freezing behavior among the groups were in the saline treated females, and only during the first 2 min but not in the following 4 min of monitoring in the test chamber (Figure 3E); however, even this apparent difference reached statistical significance only when the comparison to another group was done by t-test. More importantly, the freezing in the female control group appeared to be less than in the group treated with tatCN19o (Figure 3E); thus, even if this apparent difference was real, it would not indicate memory impairment by tatCN19o. Taken together, these data show that tatCN19o did not erase any preformed memories in our contextual fear conditioning paradigm, neither in male nor in female mice.

### tatCN19o provides potent neuroprotection in rat cortical cultures

tatCN19o is an optimized version of its parent compound, tatCN21(19). At 5 μM, tatCN21 significantly protects cultured rat hippocampal or cortical cultures from insults with glutamate even when added 1 h after the insult; however, at 1.5 μM, this neuroprotection is significantly reduced (23). Here, we decided to test neuroprotection by tatCN19o, using the same readout of neuronal cell death: release of lactate dehydrogenase (LDH) into the culture medium over 24 h after the insults. In this assay, the same level of significant neuroprotection by tatCN19o was seen after addition of 5, 2, or 0.2 μM at 1 h after the glutamate insults (Figure 4A), consistent with the improved potency of *in vitro* CaMKII inhibition by tatCN19o.

**Figure 4:**
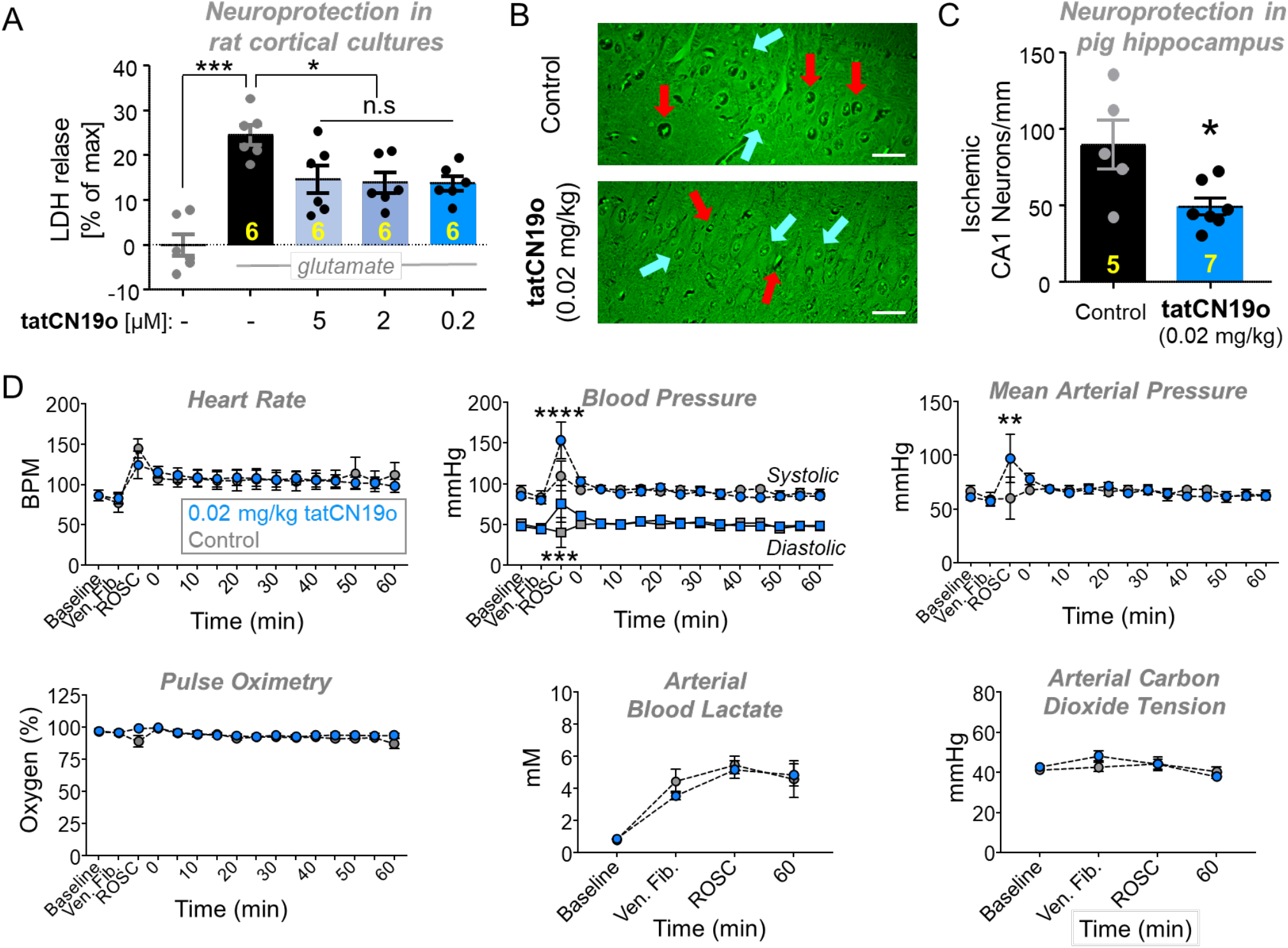
Neuroprotection with no acute cardiopulmonary effects by tatCN19o in pig.

Female pigs were injected with 0.02 mg/kg tatCN19o or saline by intravenous bolus at 30 min after resuscitation from ventricular fibrillation (i.e. return of spontaneous circulation [ROSC]), or t=0 min.

(A) First, tatCN19o was tested for protecting rat cortical cultures from neuronal cell death induced by glutamate insults (200 μM for 5 min). Cell death was assessed by release of lactate dehydrogenase (LDH) into the culture medium 24 h later; tatCN19o was added 1 h after glutamate treatment and showed significant protection at all three concentrations tested; n=6 cultures.

(B) Representative images of Fluoro-Jade B stained hippocampi from a control (top) or tatCN19o (bottom) treated animal. Examples of ischemic neurons are indicated with red arrows and healthy neurons with blue arrows. Scale bars = 100 μm.

(C) Quantification of ischemic CA1 hippocampal neurons 3 days after ventricular fibrillation. This extremely low dose achieved significant protection from ischemic injury (* p=0.0109 in two-tailed t-test; n=5 animals for saline control, n=7 animals for 0.02 mg/kg tatCN19o).

(D) Administration of tatCN19o did not appear to affect vital signs or arterial blood measures, including heart rate, systolic and diastolic blood pressure, mean arterial pressure, pulse oximetry, arterial blood lactate, and arterial carbon dioxide (n=6 for saline control, n=7 for 0.02 mg/kg tatCN19o). For all measures, no differences were seen between 0.02 mg/kg tatCN19o vs. saline treatment for any time points after injection. The only differences were observed at ROSC, the last time point before tatCN19o or saline injection (repeated measures two-way ANOVA with Bonferroni’s multiple comparisons test; p>0.9999 unless otherwise indicated: ****p<0.0001 at ROSC for systolic blood pressure, ***p=0.0009 at ROSC for diastolic blood pressure, **p=0.0011 at ROSC for mean arterial pressure, n.s. p=0.3160 at ROSC for pulse oximetry, n.s. p=0.5438 at ventricular fibrillation for arterial carbon dioxide tension; n=6 animals for saline control, n=7 animals for 0.02 mg/kg).

### tatCN19o provides potent neuroprotection also in a non-rodent species

Our previous studies showed that 0.01 mg/kg tatCN19o protects hippocampal neurons from cell death after global cerebral ischemia (GCI) induced by cardiac arrect (CA) followed by cardiopulmonary resuscitation (CPR), even when injected i.v. as single bolus at 30 min after successful CPR (18). As our behavioral studies here alleviated one of the most serious concerns regarding therapeutic use of CaMKII inhibition, we decided to further investigate the therapeutic potential of tatCN19o by testing neuroprotection in a non-rodent species. For this purpose, GCI was induced in pig by inducing ventricular fibrillations, followed by basic life support (BLS) and advanced care life support (ACLS), including defibrillation. When resuscitation was successful, 0.02 mg/kg tatCN19o was injected i.v. 30 min after return of spontaneous circulation (ROSC). This treatment with tatCN19o resulted in significant protection of hippocampal neurons, as detected by histological analysis 3 days after the injury (Figure 4B,C). Further increasing the dose to 0.2 mg/kg did not appear to further increase protective efficacy (Figure S3A); although this was tested only in two pigs, the result is consistent with our previous findings in mice. Taken together, these results show that CaMKII inhibition with tatCN19o has neuroprotective potential also in non-rodent species and suggest that maximal efficacy of tatCN19o in pig is reached at the very low dose of 0.02 mg/kg.

### No acute cardiopulmonary safety concerns for tatNC19o injection

For treatment regimens that consist of a single bolus, the highest general risk factors are acute life-threatening cardiopulmonary effects. However, no such effects were observed for the pigs that were injected with tatCN19o after defibrillation and CPR (Figures 4D and S3B,C). In these pigs, the amount of tatCN19o injected was 0.02 or 0.2 mg/kg. In mice, we additionally further escalated the dose to 10 mg/kg (i.e. to 1,000fold the therapeutic dose in this species; (18) and found no effects on heart rate or blood pressure, neither in males nor in females (Figures S4A). A common problematic drug side effect that causes cardiac complications in humans but that is not detectable in the mouse model is inhibition of hERG, a voltage-gated K^+^ channel (24,25). However, hERG binding studies *in vitro* indicated >10,000fold selectivity of tatCN19o for CaMKII over hERG and no significant binding was detected at therapeutic doses (Figures S4B), consistent also with the lack of cardiac effects in pig. Together, our results alleviate the most relevant safety concerns for tatCN19o, i.e. both the specific concerns for CaMKII inhibitors (possible effects on learning or memory) and the general concerns for any acute treatment (life-threatening acute cardiopulmonary effects).

## DISCUSSION

We showed here that short-term inhibition of CaMKII with tatCN19o did not erase preformed memories in a mouse contextual fear conditioning paradigm. Any effects on learning (i.e. memory formation) were relatively mild and very short-lived (less than 1 h). By contrast, for long-term transient interference with CaMKII signaling, our results further support the previously described memory erasure (14): Even though our expression analysis indicated significantly less expression of the T286D/T305/306AA mutant compared to the K42M tool mutant or to wildtype, the *in vivo* expression of all of these CaMKII forms was completely extinguished on day 9 after infection with the corresponding HSV-based expression vectors. Together, these results indicate that CaMKII indeed plays a role in the maintenance of memory, but also demonstrate that short-term inhibition with tatCN19o did not cause any retrograde amnesia or long-term impact on learning that could be contraindications for its therapeutic use in neuroprotection.

The most convincing study to date to suggest a role of CaMKII in the maintenance of hippocampus-dependent memory utilized transient virus-mediated expression of the kinase-dead K42M mutant in rat hippocampus after the rats learned a place avoidance task (14). The viral vector was based on HSV, which is known to cause a transient protein expression that subsides within the weeks after injection. For testing the effects on memory maintenance, it is crucial that the CaMKII mutant is not longer expressed on the day of the memory test; otherwise, any apparent reduction in memory may be due to acute effects on memory recall rather than persistent effects on memory maintenance beyond the transient expression of the mutant. However, whereas expression of the CaMKII T286D/T305/306AV mutant was shown to be eliminated on the day of testing, a similar elimination of expression of CaMKII wildtype or the more important K42M mutant was not shown (14). Whereas our results now show that the K42M mutant is significantly more highly expressed than the T286D/T305/306AV mutant, they also show that after HSV-mediated expression, the expression is transient and eliminated on day 9 (the day of testing) not only for the T286D/T305/306AV mutant but also for the higher expressing CaMKII wildtype and K42M mutant. This indicates that prolonged expression of the K42M can indeed reverse preformed memories. By contrast, pharmacological short-term inhibition of CaMKII with tatCN19o did not cause reversal of memory in our mouse contextual fear conditioning paradigm.

Even though tatCN19o did not erase pre-formed memories, it mildly decreased memory formation, i.e. learning. Similar results have been obtained previously with tatCN21, the less potent parent compound of tatCN19o (26). Importantly, our results show that the mild impairment in learning was very transient and completely reversed within one hour. Even transient learning impairment could be a contraindication for therapeutic application in some conditions. However, for global cerebral ischemia or stroke, which are the currently the main potential target conditions for treatment with tatCN19o (18,23), such transient learning impairment would be highly acceptable even if they were longer lasting than observed here, especially since the treatment is with a single acute bolus of the drug. Additionally, the very short duration of the learning impairment could enable even chronic treatments of some conditions. Such conditions could include Alzheimer’s disease, because CaMKII mediates not only normal LTP but also the LTP impairment caused by Aβ (27,28), one of the pathological agents associated with Alzheimer’s disease. However, this would require that the restoration of Aβ-related LTP impairments by CaMKII inhibition is significantly longer lasting than the acute transient LTP impairment that is directly caused by the treatment. By contrast, for global cerebral ischemia, our results directly support further development of tatCN19 towards a human therapy: In addition to alleviating primary specific concerns regarding effects on memory, our results also alleviate secondary general concerns regarding potential cardiopulmonary risks, and they demonstrate efficacy in a non-rodent species in a highly clinically relevant injury and treatment model (ventricular fibrillation followed by BLS and ACLS).

Taken together, our results support that CaMKII signaling is indeed involved in the maintenance of memory, but that at least short-term inhibition of CaMKII does not cause a reversal of memory that could be a contraindication for therapeutic applications.

## EXPERIMENTAL PROCEDURES

### Material availability

Requests for resources, reagents, or questions about methods should be directed to K. Ulrich Bayer. This study did not generate new unique reagents.

### Animal and cell culture models

The experiment with HSV *in vivo* injection were conducted in accordance with Brandeis University and the Institutional Animal Care and Use Committee (IACUC) approved protocols, and utilized adult male Long-Evans rats (2-3 months old) that were housed 1-2 per cage in a temperature-controlled room with a 12-12 h light/dark cycle. All other animal treatments were approved by the University of Colorado Institutional Animal Care and Use Committee, in accordance with NIH guidelines, and was done as described previously (29). Pregnant Sprague-Dawley rats and adult male C57BL/6 mice, 8 to 12 weeks old, were supplied by Charles River Labs. Animals are housed at the Animal Resource Center at the University of Colorado Anschutz Medical Campus (Aurora, CO) and are regularly monitored with respect to general health, cage changes, and overcrowding. Pigs were housed, and experimentation took place in the animal care facility at the University of Colorado Anschutz Medical Campus. Specific pathogen free, female Yorkshire cross swine (12-14 weeks old) were purchased from Oak Hill Genetics.

### Material and DNA constructs

Material was obtained from Sigma, unless noted otherwise. Co-expression of SYFP2-CaMKII and mTurquoise2 (referred to as YFP and mTurquois) was driven by an IRES vector that we have used previously (30).

### Western analysis

Before undergoing SDS-PAGE, samples were boiled in Laemmli sample buffer for 5 min at 95° C. Proteins were separated in a resolving phase polymerized from 9% acrylamide, then transferred to a polyvinylidene difluoride membrane at 24 V for 1 h at 4°C. Membranes were blocked in 5% milk and incubated with anti-GFP (1:1000) followed by Amersham ECL goat anti-rabbit horseradish peroxidase conjugate 1:5000 (Bio-Rad). Blots were developed using chemiluminescence (Super Signal West Femto, Thermo-Fisher), imaged using the Chemi-Imager 4400 system (Alpha-Innotech), and analyzed by densitometry (ImageJ). Phospho-signal was corrected to total protein. Relative band intensity for YFP-CaMKII was normalized as a ratio of mTurquoise intensity on the same blot.

### Proteasomal inhibition with MG-132

MG-132 was solubilized in DMSO. HEK cells or neurons were treated with either DMSO or 50 μM MG-132 4 hours prior to harvest or imaging.

### Protein extracts

Whole homogenates of HEK cells were made in homogenization buffer including 50 mM PIPES pH 7.2, 10% glycerol, protease inhibitor cocktail and 1 mM DTT. The homogenates were cleared by centrifugation at 14,000 RPM at 4°C for 20 min. For all expression experiments, cells were harvested 24 h after transfection.

### Primary hippocampal culture preparation

To prepare dissociated neuronal cultures, cortex or hippocampi were dissected from mixed sex rat pups (P0), dissociated in papain for 1 h, and plated at 100,000 cells/mL on glass coverslips for imaging and 500,000 cells/mL on culture dishes for biochemistry. At DIV 12-14, hippocampal neurons were transfected with 1 μg total cDNA per well using Lipofectamine 2000 (Invitrogen), then imaged or treated and fixed 2-3 days later. At DIV 14, cortical neurons were subjected to glutamate insult and later assessed for cell death.

### Image analysis in HEK cells or neurons

Regions of interest (ROIs) were hand drawn around the cytoplasm of HEK cells or the cell body of hippocampal neurons. Both YFP and mTurquoise intensity were quantified in this region, and the final values are expressed as the ratio of YFP to mTurquoise.

### Excitotoxic cell death in neuronal cultures

DIV 14 cortical neurons were subjected to water control or glutamate insult (200 μM glutamate for 5 min). Prior to the insult, half of the media was removed (and saved for later use as conditioned media) from each well and replenished with fresh feeding media. After the insult, media was exchanged with a 1:1 mix of fresh and conditioned media and neurons were returned to 37°C and 5% CO2 for 20-24 h. Media samples to be tested for cell death were collected 1 h before or 24 h after glutamate insult, as indicated. Cell death was then assessed as described previously (23,31) by measuring the lactate dehydrogenase (LDH) released from the cells into the media using a Pierce LDH Cytotoxicity Assay Kit (Thermo Scientific).

### Contextual Fear Conditioning

The mice were allowed to acclimate for at least one week and handled daily at least 3 days before behavioral testing. On the training day, the mice are placed into conditioning box with a metal grid on the floor (MedAssociates). Over a 6 min period in the conditioning box, the animals received 2 separate foot shocks (2 s, 0.7 mA, 2 min interval). The mice are then placed back into their home cage. 24 h later, on the testing day, the mice are placed into the same box for a 6 min trial without any foot shocks. Movement was measured by a video monitoring system (FreezeScan 2.0) during both days of testing and freezing behavior was calculated as the percentage of time the mice were immobile.

### Global cerebral ischemia in pig by ventricular fibrillation

Global cerebral ischemia (GCI) in pig was induced by delivering a 60Hz, 100 mA electric current across the thorax resulting ventricular fibrillation (VF) as confirmed by ECG. After 4 minutes of VF, basic life support (BLS) was initiated according to guidelines from the American Heart Association (AHA). Longer periods of VF (6 and 10 minutes) were attempted in an effort to maximize injury to the hippocampus, but resulted in high mortality (50% and 90%, respectively); thus we chose 4 minutes of VF which resulted in a mortality of 10%. Briefly chest compressions and breaths were delivered at a rate of 30:2 compressions to ventilations to maintain an end tidal CO_2_ (ETCO_2_) of 15-20 mmHg. After 2 minutes of BLS, advanced cardiac life support (ACLS) following AHA guidelines commenced. During ACLS, compressions are delivered at a continuous rate and ventilations are delivered via ventilator at a set respiratory rate of 18 breaths per minute and a tidal volume of approximately 300 ml (8 ml/kg). After return of spontaneous circulation (ROSC; indicated by an ETCO_2_ of 30-40 mmHg and return of heart rate and rhythm) chest compressions and ventilator support was stopped. After 3 days, animals were euthanized with sodium pentobarbital, according to AVMA guidelines, and brains collected for histological analysis of hippocampal neuronal cell death.

The procedures were performed under anesthesia, induced with intramuscular administration of 10-20 mg/kg ketamine and 3-5% isoflurane via nosecone. Animals were intubated with a cuffed 8.0 mm endotracheal tube, and peripheral venous access obtained. Sedation was maintained throughout the experiment with 1-3% isoflurane. The external jugular vein and femoral artery were accessed using ultrasound guided percutaneous micropuncture. Respiratory parameters, pulse oximetry, body temperature, invasive blood pressure, and electrocardiogram (ECG) were monitored throughout the experiment. Following resuscitation and prior to waking from anesthesia, pigs received a subcutaneous injection of 0.12 mg/kg of buprenorphine SR for pain relief and were allowed to recover in their pens. Following recovery, pigs were observed twice daily. Animals with the inability to stand/ambulate after recovering from anesthesia, suffering from weakness resulting in the inability to eat or drink for greater than 24 hours, showing signs of central nervous system depression resulting in seizures, paralysis, or suffering from pain unresponsive to analgesia are euthanized prior to the end of the study. Animals with a sustained mean arterial pressure for 10 continuous minutes during resuscitation were also euthanized.

### Histological analysis of neuronal cell death

Fluoro-Jade B (FJB) staining was performed to evaluate neuronal injury in the dorsal hippocampus. Coronal 6 μm paraffin sections were deparaffinized and rehydrated. Slides were immersed in 1% NaOH in 80% alcohol solution for 5 min, then immersed in 70% alcohol and distilled water for 2 min. Slides were blocked in 0.06% potassium permanganate for 20 min before being stained in 0.004% FJB solution in 0.1% acetic acid for 20 min. Slides were rinsed with distilled water, dried on a slide warmer for 10 min, and dipped in xylene and mounted on coverslips with DPX mountant. Images were acquired on a epifluorescent microscope using a 20x objective. 3 ROI’s in 3 sections were obtained per animal. Image analysis was performed by a blinded investigator. Ischemic neurons were characterized by a condensed FJB positive nucleus that did not have chromatin strands or healthy cytoplasm surrounding them.

### hERG channel interaction in vitro

tatCN19o inhibition of [^3^H]dofetilide binding to hERG channel protein was used to assess its potential cardiotoxicity. hERG channel protein was stably expressed in HEK-293 cells purchased from Millipore (Cat. No. CYL3006; Billerica, MA). Assay methods have been previously described (32–34). Frozen HEK-293 cells were thawed at 37°C and placed for 4-8 h in 20 ml of complete media (minimum essential medium supplemented with 10% fetal bovine serum, 1% nonessential amino acids, and 400 mg/ml geneticin) in T-75 cm^2^ flasks (Becton Dickinson and Company, Franklin Lakes, NJ) in a humidified atmosphere (5% CO_2_, 37°C). Media was replaced with 20 ml fresh media after 4-8 h and every 48 h subsequently. For routine cell passages (every 6 days), media was removed, cells rinsed with 2 ml of phosphate-buffered saline (137 mM sodium chloride, 2.7 mM potassium chloride, 10 mM disodium hydrogen phosphate, 2 mM potassium dihydrogen phosphate), and Hank’s balanced salt solution containing trypsin (0.5 g/l, porcine trypsin) and EDTA (0.5 mM) was added. For cell dissociation, flasks were placed in a 37°C incubator for 2-5 min. Fresh complete media (5 ml) was added to the cell suspensions, and cells were seeded onto new flasks at 2–3 × 10^6^ cells/flask. On the last passage prior to membrane preparation, cells were seeded onto culture dishes (150 mm x 25 mm) at 2.5 × 10^6^ cells/dish and incubated (5% CO_2_ at 37°C) for 40-48 h. Media was removed and dishes were rinsed twice with 30°C Hanks’ balanced salt solution (13 ml). A solution of ice-cold 0.32 M sucrose with 5 mM sodium bicarbonate (20 ml, pH 7.4) was added to each dish on ice. Cells were scraped gently off the dishes and homogenized (30 s) on ice with a Teflon pestle (0.003 in) using a Maximal Digital homogenizer (280 rpm). Homogenates were centrifuged (300g and 800g, 4 min each, 4°C). Pellets were resuspended in 9 ml ice-cold MilliQ water. Osmolarity was restored by addition of 1 ml of 500 mM Tris buffer (pH 7.4). Samples were centrifuged (20,000g, 30 min, 4°C). Pellets were resuspended in 2 ml assay buffer (50 mM Tris, 10 mM potassium chloride, and 1 mM magnesium chloride, pH 7.4, 4°C). Aliquots of membrane suspension were stored at −80°C. For the [^3^H]dofetilide binding assay, membrane suspension was thawed and protein content determined using a Bradford protein assay (Bio-Rad Laboratories, Inc., Hercules, CA), with bovine albumin (Sigma-Aldrich Corporation) as the standard. Duplicate tubes were prepared containing membrane suspension (5 mg/100 ml), one of a range of concentrations of tatCN19o (final concentrations 0, 0.1 nM to 0.1 mM in 25 ml) or amitriptyline (0, 0.1 nM–0.1 mM; positive control), assay buffer (150 ml), and [^3^H]dofetilide (5 nM in 25 ml) for a final assay volume of 250 ml. Amitriptyline (1 mM) was used to determine nonspecific binding (35,36). Samples were incubated for 1 h at room temperature. Reactions were stopped by rapid filtration through Whatman GF/B Glass microfiber filters presoaked in 0.5% PEI for 1 h at 4°C. Filters were washed 3 times with 1 ml ice-cold assay buffer. Radioactivity retained on the filters was determined by liquid scintillation spectrometry (TRI-CARB 2100 TR Packard scintillation counter; Packard BioScience Company, Meriden, CT).

### Quantification and statistical analysis

All data are shown as mean ± SEM. Statistical significance is indicated in the figure legends. Statistics were performed using Prism (GraphPad) software. Imaging experiments were obtained and analyzed using SlideBook 6.0 software. Western blots were analyzed using ImageJ (NIH). All data were tested for their ability to meet parametric conditions, as evaluated by a Shapiro-Wilk test for normal distribution and a Brown-Forsythe test (3 or more groups) or an F-test (2 groups) to determine equal variance. All comparisons between two groups met parametric criteria, and independent samples were analyzed using unpaired, two-tailed Student’s t-tests. Comparisons between three or more groups meeting parametric criteria were done by one-way ANOVA with specific post-hoc analysis indicated in figure legends. Comparisons between three or more groups with two independent variables were assessed by two-way ANOVA with Bonferroni post-hoc test to determine whether there is an interaction and/or main effect between the variables.

## ACKNOWLEDGEMENTS

We thank Dr. Steven Coultrap and Ms. Janna Mize-Berge for help with mouse colony maintenance.

## AUTHOR CONTRIBUTIONS

N.L.R., C.N.B., T.B.H., T.R., J.E.O, J.E.T., L.P.D., O.R.B., and N.Q. performed experiments and analyses; N.L.R., C.N.B., J.E.L., P.S.H, V.S.B., and K.U.B. conceived this study; K.U.B. wrote the initial draft and all authors contributed to the final manuscript.

## FUNDING AND ADDITIONAL INFORMATION

This work was funded by National Institutes of Health grants T32 GM007635 (supporting C.N.B.), T32 AG000279 (supporting N.L.R.), F31 AG069458 (to N.L.R.), F31 NS129254 (to C.N.B.), F32 AG066536 (to O.R.B.), R01 AG067713, R01 NS110383, R01 NS081248 (to K.U.B.), and R01 NS118786 (to K.U.B. and P.S.H.).

