## Supplemental Figures 1-4 for "Short-term CaMKII inhibition with tatCN19o does not erase pre-formed memory and is neuroprotective in non-rodents"

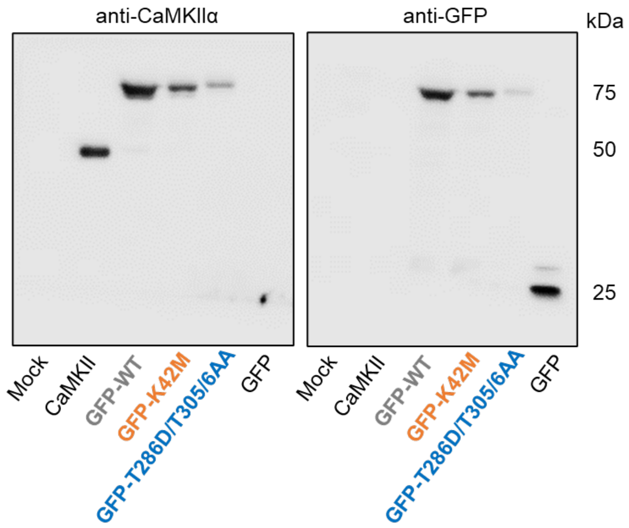


**Supplemental Figure S1: No apparent degradation products for GFP-CaMKII wildtype, K42M, or T286D/T305/6AA CaMKII in HEK cells.** Western analysis of lysates from HEK cells transfected with nothing (Mock), untagged CaMKII, GFP-CaMKII wildtype (WT) and tool mutants (K42M and T286D/T305/6AA), and GFP alone. Blots were probed with an anti-CaMKII antibody, stripped, and reprobed with an anti-GFP antibody. No degradation products were detected for any of the constructs with either antibody.

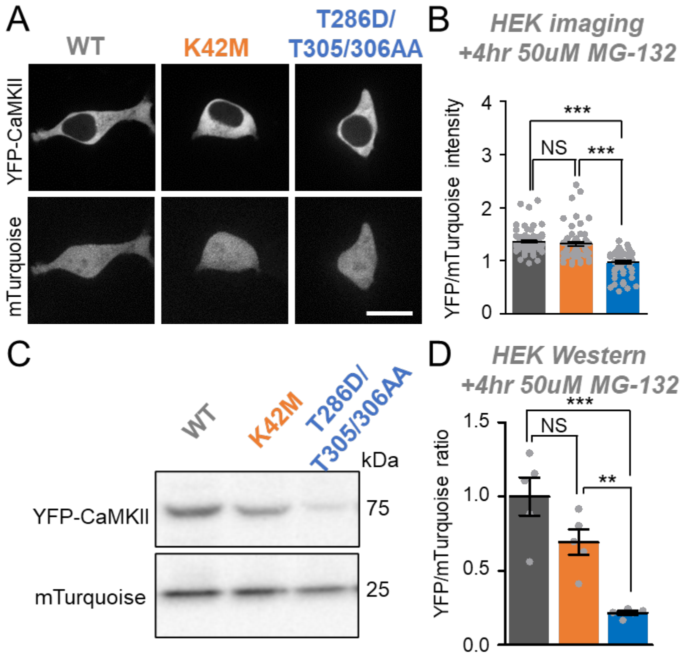


**Supplemental Figure S2: Acute short-term proteasome inhibition does not restore CaMKII expression.** Error bars indicate SEM in all panels. **P<0.01; ***P<0.001; NS, not significant by one-way ANOVA with post-hoc by Tukey's test.

**(**A) An mTurquoise-IRES-YFP-CaMKII vector was transfected into HEK cells for 24h. 50 μM MG-132 was added for the last 4 hours and cells were analyzed for mTurquoise and YFP-CaMKII expression by confocal imaging. Scale bar=10μm. (**B**) Quantification from imaging experiments. (**C**) Representative western blots showing mTurquoise and YFP-CaMKII expression after 4 hours of 50 μM MG-132 treatment with (**D**) densitometric quantification.

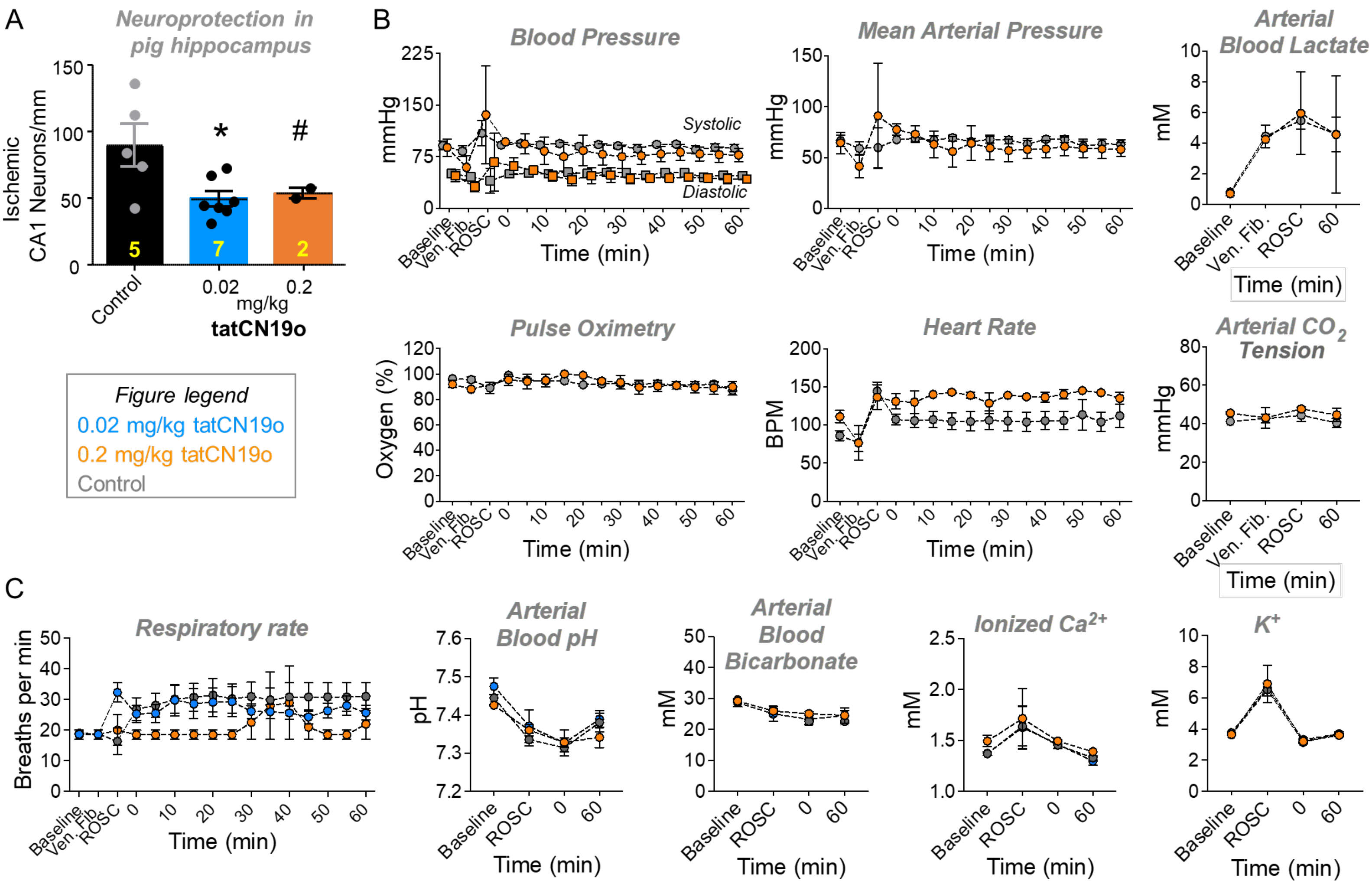


**Supplemental Figure S3: Maximal efficacy with no acute cardiopulmonary effects by tatCN19o in pig.** Female pigs were injected with 0.02 or 0.2 mg/kg tatCN19o (or saline) by intravenous bolus 30 min after resuscitation from ventricular fibrillation (i.e. return of spontaneous circulation [ROSC]), or t=0 min. (**A**) The low dose of 0.02 mg/kg appears to achieve maximal protection from ischemic injury in CA1 hippocampal neurons, as further increasing the to 0.2 mg/kg did not appear to further increase efficacy of protection (although only 2 animals were tested at this higher dose). (**B**) 0.2 mg/kg tatCN19o did not appear to affect vital signs or arterial blood measures, including heart rate, mean arterial pressure, systolic and diastolic blood pressure, pulse oximetry, arterial blood lactate, and arterial carbon dioxide. (**C**) Additional measures for 0.02 and 0.2 mg/kg tatCN19o included respiratory rate, arterial blood pH, arterial blood bicarbonate, ionized calcium, and potassium. No differences were seen between 0.02 mg/kg tatCN19o vs. saline treatment for any time points after injection. The only differences were observed before tatCN19o or saline injection (repeated measures two-way ANOVA with Bonferroni's multiple comparisons test; $p>0.9999$ unless otherwise indicated: n.s. $p=0.8419$ at ROSC for respiratory rate; n=6 animals for saline control, n=7 animals for 0.02 mg/kg).

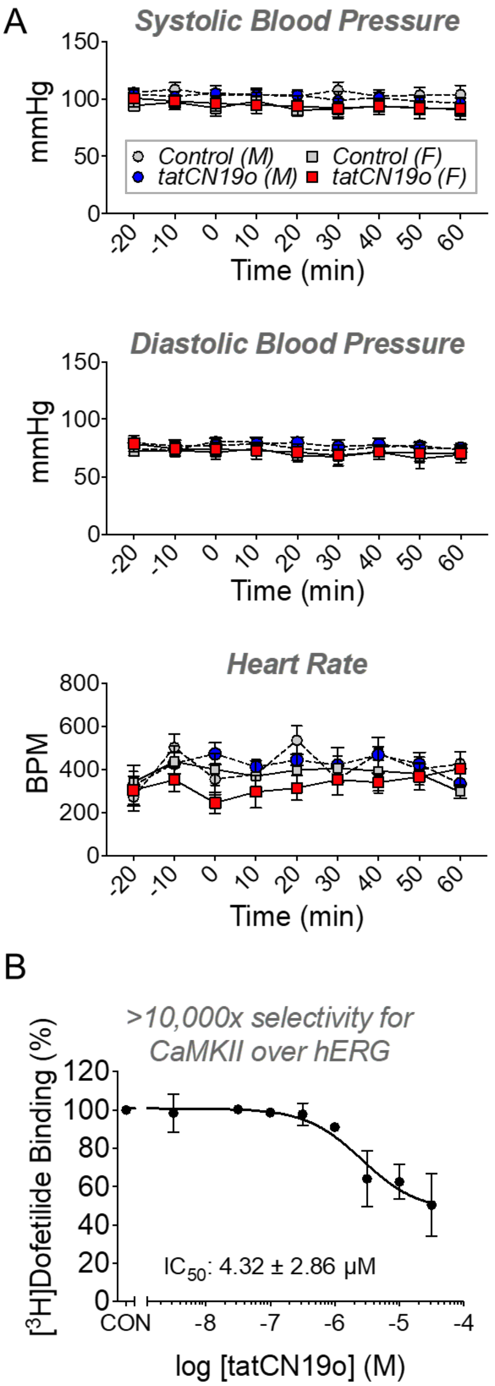


**Supplemental Figure S4: No acute cardiac effects by tatCN19o in mice and *in vitro*.** (**A**) Male and female mice were injected with 10 mg/kg tatCN19o (1,000fold the therapeutic dose) by intravenous bolus at t=0, or 30 min after CPR recovery from cardiac arrest. No acute effects on systolic or diastolic blood pressure (n=6 males, n=6 females per treatment group; genders were analyzed separately by repeated measures two-way ANOVA with Bonferroni's multiple comparisons test; p>0.9999 for all post-hoc comparisons). The same was true for heart rate (n=3 males per treatment, n=4 females per treatment; repeated measures two-way ANOVA with Bonferroni's multiple comparisons test; p>0.9999 unless otherwise indicated: NS p=0.6425 at t=0 for females). (**B**) Preliminary *in vitro* hERG assay showed that at therapeutic doses (50 nM initial blood stream concentration), no binding to the hERG channel was detected at all (n=3 dose-response binding assays). With ~4.3 µM, the $IC_{50}$ for hERG binding was >10,000fold higher compared to the $IC_{50}$ for CaMKII inhibition of <0.4 nM.
